# Dicer accumulates in cytoplasmic foci upon alphavirus infection and plays a proviral role in *Myotis myotis* bat cells

**DOI:** 10.1101/2025.07.10.664130

**Authors:** Léa Gaucherand, Hugo Marie, Julie Cremaschi, Sébastien Pfeffer

## Abstract

Bats are reservoirs for many viruses that frequently cause epidemics in humans and animals. It is thus critical to better understand their immune system and mechanisms of antiviral immunity. Despite an increasing number of studies, much is still unknown about the molecular mechanisms that govern bat-virus interactions, especially given the large diversity of bat species. Dicer is a conserved ribonuclease with multiple activities that can modulate antiviral immunity, including the detection of viral RNA as part of the RNA interference (RNAi) pathway, the maturation of micro RNAs, and the direct inhibition of innate immunity in mouse and human cells. In view of these complex activities of Dicer, we tested its antiviral activity in *Myotis myotis* nasal epithelial cells. Surprisingly, we did not see strong evidence of RNAi in these cells, but instead saw a proviral effect of Dicer for two alphaviruses, Sindbis and Semliki forest virus. We also observed a striking relocalization of Dicer to cytoplasmic foci upon infection with these viruses, which did not occur in the several human cell lines we tested. These foci contained dsRNA and viral plus strand RNA, suggesting that they are sites of viral replication. Finally, we found that factors specific to *M. myotis* cells are needed for Dicer relocalization. Overall, we propose that Dicer can play different roles in different bat species and/or cell types, and is being repurposed by alphaviruses to promote viral replication.

## Introduction

Bats are unique mammals in many aspects, such as their ability to fly and their high longevity relative to their body size (Gonzalez & Banerjee, 2022). They belong to the order Chiroptera, which can be divided into two suborders that diverged over 50 million years ago (Lei & Dong, 2016). More than 1400 different species of bats have been identified so far, accounting for 20% of all mammals (Burgin *et al*, 2018). Importantly, bats are reservoirs for many viruses that can spillover into humans, either directly or through an intermediate host, such as Marburg virus, Hendra and Nipah virus, Middle East respiratory syndrome coronavirus (MERS-CoV), severe acute respiratory syndrome coronavirus (SARS-CoV) and likely SARS-CoV-2 (Halpin *et al*, 2000; Li *et al*, 2005; Memish *et al*, 2013; Moratelli & Calisher, 2015; Chua *et al*, 2002; Yob *et al*, 2001; Towner *et al*, 2007). Interestingly, while these viruses can be very pathogenic to humans and other animals, most of the studied bats so far show little to no clinical symptoms, whether they were naturally or experimentally infected (Gonzalez & Banerjee, 2022; Pei *et al*, 2024). An increasing number of studies are highlighting the role of the bat innate immune system in this apparent tolerance to viruses.

The type I and III interferon (IFN) pathways are the main first lines of defense against viruses in mammals. In this pathway, sensing of viral nucleic acids by pattern recognition receptors (PRRs) leads to the expression and secretion of type I and III IFN cytokines. These cytokines are then sensed by cells in an autocrine and paracrine fashion, leading to the expression of interferon stimulated genes (ISGs) that can actively fight the infection (Pichlmair & Reis e Sousa, 2007). The IFN pathway is present and active in bats, but differences have been reported compared to humans, as well as between different bat species. These include a dampening of the response to DNA sensing, as well as differences in the baseline expression and temporal regulation of some IFNs and ISGs (Irving *et al*, 2021; Banerjee *et al*, 2020; Clayton & Munir, 2020). In contrast, invertebrates and plants mostly rely on the RNA interference (RNAi) pathway to fight viral infections. In antiviral RNAi, viral double-stranded (ds)RNAs are sensed and cleaved into small interfering (si)RNAs by a Dicer or Dicer-like ribonuclease, then loaded into a protein of the Argonaute family to target and degrade complementary viral RNAs (Guo *et al*, 2019). Interestingly, two recent studies have reported enhanced antiviral activity of Dicer through the RNAi pathway in the *Pteropus alecto* kidney (PaKi) cell line infected with SINV, an NS1 deletion mutant strain of infuenza A virus and a B2 deletion mutant strain of Nodamura virus, and in the *Tadarida brasiliensis* lung (TBLU) cell line infected with SARS-CoV-2 (Dai *et al*, 2024; Owolabi *et al*, 2025). The first study also proposed that viral dsRNA cleavage by Dicer limits the recognition of dsRNA by PRRs and thus the activation of the IFN response in *P. alecto*, thereby suggesting a role for Dicer in bat viral tolerance (Dai *et al*, 2024).

Dicer is well known for its essential role in the maturation of most micro (mi)RNAs, key regulators of gene expression (Bartel, 2018). In mammals, the same Dicer protein can generate both miRNAs and siRNAs of 22 nucleotides (nt) thanks to its regulatory cofactors transactivation response element RNA-binding protein (TRBP) and Protein ACTivator of the interferon-induced protein kinase (PACT) (Lee *et al*, 2013). However, the antiviral activity of Dicer through the RNAi pathway is controversial and appears to be cell-type and context-dependent in human and mouse cells (Cullen *et al*, 2013). Indeed, Dicer isoforms with increased RNAi activity have been identified in human and mouse stem cells (Poirier *et al*, 2021), and in mouse oocytes (Flemr *et al*, 2013). While the former appears to have some antiviral properties against viruses such as Zika and SARS-CoV-2 (Poirier *et al*, 2021), the mouse oocyte specific one does not confer an antiviral phenotype when artificially expressed ubiquitously (Kulmann *et al*, 2025). One important reason for the reduced antiviral activity of full-length Dicer is the existence of a regulatory crosstalk between the IFN and RNAi pathways (Baldaccini & Pfeffer, 2021; Gaucherand *et al*, 2024; Maillard *et al*, 2019). Human Dicer is inhibited by the PRR LGP2 (laboratory of genetics and physiology 2 / DexH box polypeptide 58) (van der Veen *et al*, 2018). Moreover, inactivating the IFN pathway in differentiated mouse cells unmasks long dsRNA processing by Dicer (Maillard *et al*, 2016). Conversely, Dicer represses IFN and the ISG protein kinase R (PKR) activity in mouse embryonic stem cells transfected with double stranded RNA (Gurung *et al*, 2021). In addition, we recently showed that human Dicer interacts with PKR *via* its helicase domain and regulates innate immune activation during Sindbis virus (SINV) infection (Montavon *et al*, 2021; Baldaccini *et al*, 2024).

Overall, Dicer could modulate antiviral activity through its role in the RNAi pathway, but also by maturing miRNAs that can affect viral infection, by directly regulating innate immune activation, and potentially through other still unknown mechanisms. How these multiple complex activities of Dicer could contribute to antiviral activity and viral tolerance in different bat species is still poorly understood. This prompted us to investigate the antiviral activity of Dicer in the greater mouse-eared bat (*Myotis myotis*), a European microbat from the *Vespertilionidae* family. Unexpectedly, expressing *M. myotis* Dicer in human Dicer knock out cells did not protect from viral infection. Instead, we saw a proviral role of Dicer in *M. myotis* nasal epithelial cells upon infection with two different alphaviruses, with no clear evidence of antiviral RNAi activity. Interestingly, Dicer relocalizes to large cytoplasmic foci upon infection with these viruses in *M. myotis* nasal epithelial cells but not in human cells. We find that these foci are likely viral factories and that *M. myotis* cell*-*specific factors are required for Dicer relocalization. Overall, we propose that the activity of bat Dicer is context-dependent and could be used by some viruses like SINV in *M. myotis* cells to promote viral replication.

## Results

### Expressing *M. myotis* Dicer in NoDice cells does not protect from SINV and VSV infection

Since *P. alecto* and *T. brasiliensis* Dicer were found to be antiviral against a few viruses (Dai *et al*, 2024; Owolabi *et al*, 2025), we tested the antiviral activity of *M. myotis* Dicer. We cloned Dicer from *M. myotis* cells (hereafter referred to as mmDicer) and expressed it with an HA tag in human embryonic kidney (HEK) 293T cells knock-out for Dicer (NoDice cells (Bogerd *et al*, 2014)). As controls, we also expressed an HA-tagged catalytically inactive mutant (amino acid changes E1562A and E1819A) version of *M. myotis* Dicer (hereafter referred to as mmDicer CM), HA-tagged human Dicer (hDicer), or an empty vector (Empty). Efficient expression of the different Dicer constructs was checked by western blot (**Fig. 1A**). Expressing mmDicer but not mmDicer CM also rescued maturation of the human miRNA miR-16 (**Fig. 1B**). These results suggested that mmDicer is indeed expressed and active in NoDice cells. We then tested its antiviral activity against a SINV that expresses GFP (SINV-GFP) (López *et al*, 2020). Expressing mmDicer did not lead to a significant decrease in SINV-GFP titers, RNA and protein levels compared to the empty vector control, hDicer or mmDicer CM (**Fig. 1C-E**), suggesting that mmDicer is not particularly antiviral against SINV in this context. We also tested a different virus, the rhabdovirus vesicular stomatitis virus expressing GFP (VSV-GFP), and again saw no antiviral activity of mmDicer (**Fig. 1F**). Overall, these results suggest that expressing mmDicer in a human cell line does not protect against SINV and VSV infection.

**Figure 1:**
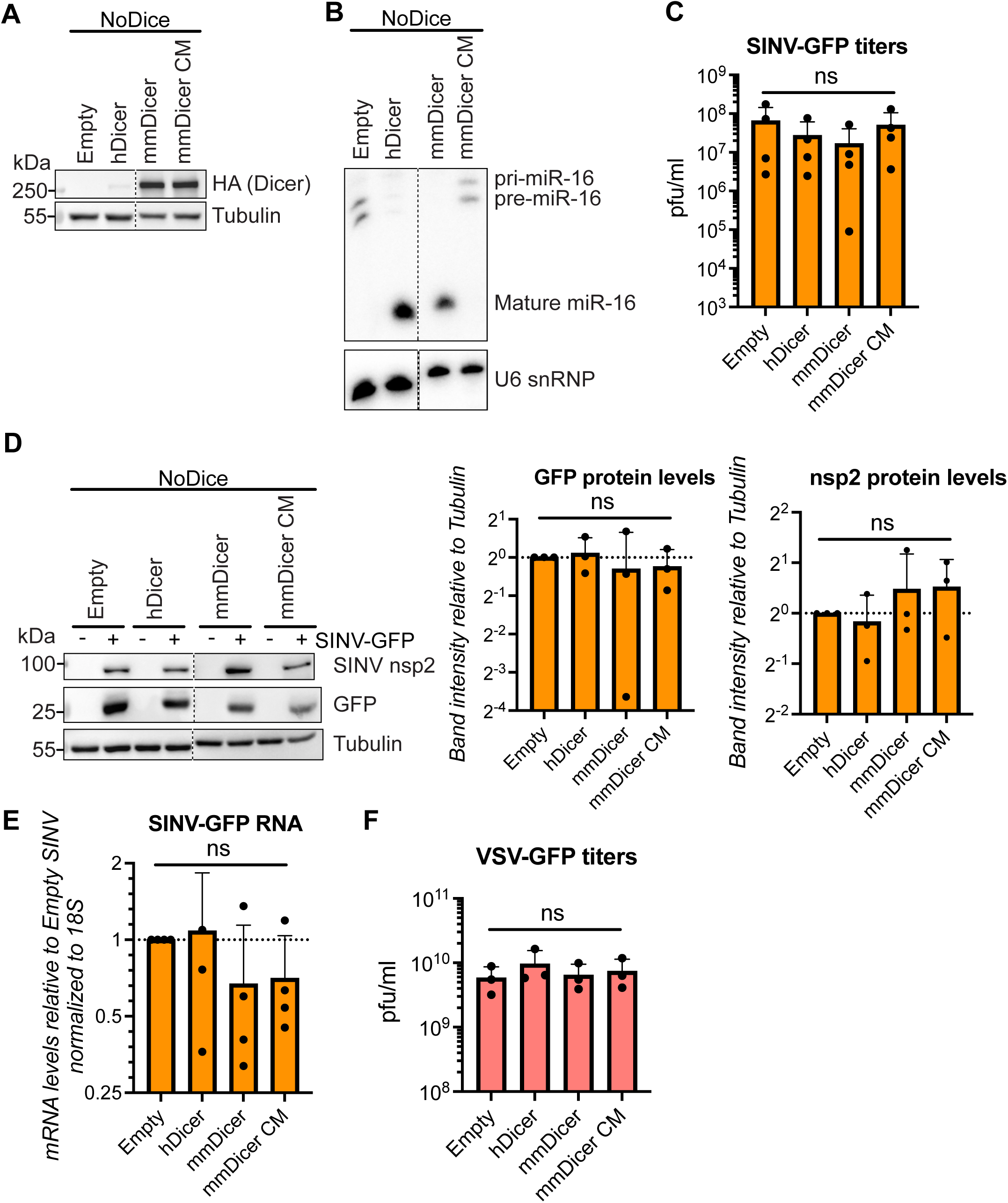
Expressing *M. myotis* Dicer in human NoDice cells does not protect from SINV and VSV infection. **(A)** Expression of HA-tagged Dicer was analyzed by western blot for NoDice cells expressing HA-tagged hDicer, mmDicer, mmDicer CM or an empty vector control. α-Tubulin was used as loading control. Images are representative of 3 independent experiments **(B)** Expression of miR-16 was analyzed by northern blot for NoDice cells expressing HA-tagged hDicer, mmDicer, mmDicer CM or an empty vector control. A probe for U6 snRNP was used as loading control. Images are representative of 2 independent experiments. **(C-F)** NoDice cells expressing HA-tagged hDicer, mmDicer, mmDicer CM or an empty vector control were infected with SINV-GFP **(C-E)** or VSV-GFP **(F)** at MOI 0.02 for 24h. **(C, F)** Supernatants were collected and viral titers were quantified by plaque assay. Means +/- standard deviation are plotted for 4 **(C)** or 3 **(F)** independent experiments. **(D)** Protein lysates were collected and analyzed by western blot using antibodies for SINV nsp2 and GFP. α-Tubulin was used as loading control. n = 3 independent experiments. Quantification of GFP or nsp2 band intensities relative to Tubulin band intensities were quantified with ImageJ and the mean +/- standard deviation was plotted relative to empty vector control for all three independent experiments. **(E)** RNAs were purified and levels of SINV RNA were quantified by qRT-PCR. Means +/- standard deviation normalized to 18S are plotted relative to empty vector control for 4 independent experiments. For all graphs, p values were calculated using standard one-way ANOVA with Dunnett’s multiple comparison test. ns: not significant.

### Dicer is proviral for two alphaviruses in *M. myotis* nasal epithelial cells

To determine whether mmDicer displays antiviral activity in *M. myotis* cells, we used a pool of multiple siRNAs at low concentrations to knock down Dicer in a SV40 transformed *M. myotis* nasal epithelial cell line (He *et al*, 2014b). This technique allows for efficient knock down (**Fig. 2A**) while limiting off-target effects. We then infected these cells with SINV-GFP and VSV-GFP. Interestingly, we saw that knocking down Dicer actually reduced SINV-GFP replication, as seen by a decrease in GFP+ cells over time (**Fig. 2B**). At 24h, we also saw a decrease in SINV-GFP titers, in GFP protein levels and in SINV RNA levels (**Fig. 2C-E**). To test whether the proviral effect of Dicer could be recapitulated with another alphavirus, we infected our *M. myotis* cell line with Semliki forest virus (SFV). Similar to SINV infection, knocking down Dicer decreased SFV titers and RNA levels at 24 hours post infection (**Fig. 2F-G**). In contrast, Dicer knock down had a proviral effect against VSV-GFP infection at the RNA and protein levels, but no difference in titers was found (**Fig. 2H-J**). Overall, our results suggest that Dicer plays a proviral role for SINV and SFV but not VSV infection in *M. myotis* nasal epithelial cells.

**Figure 2:**
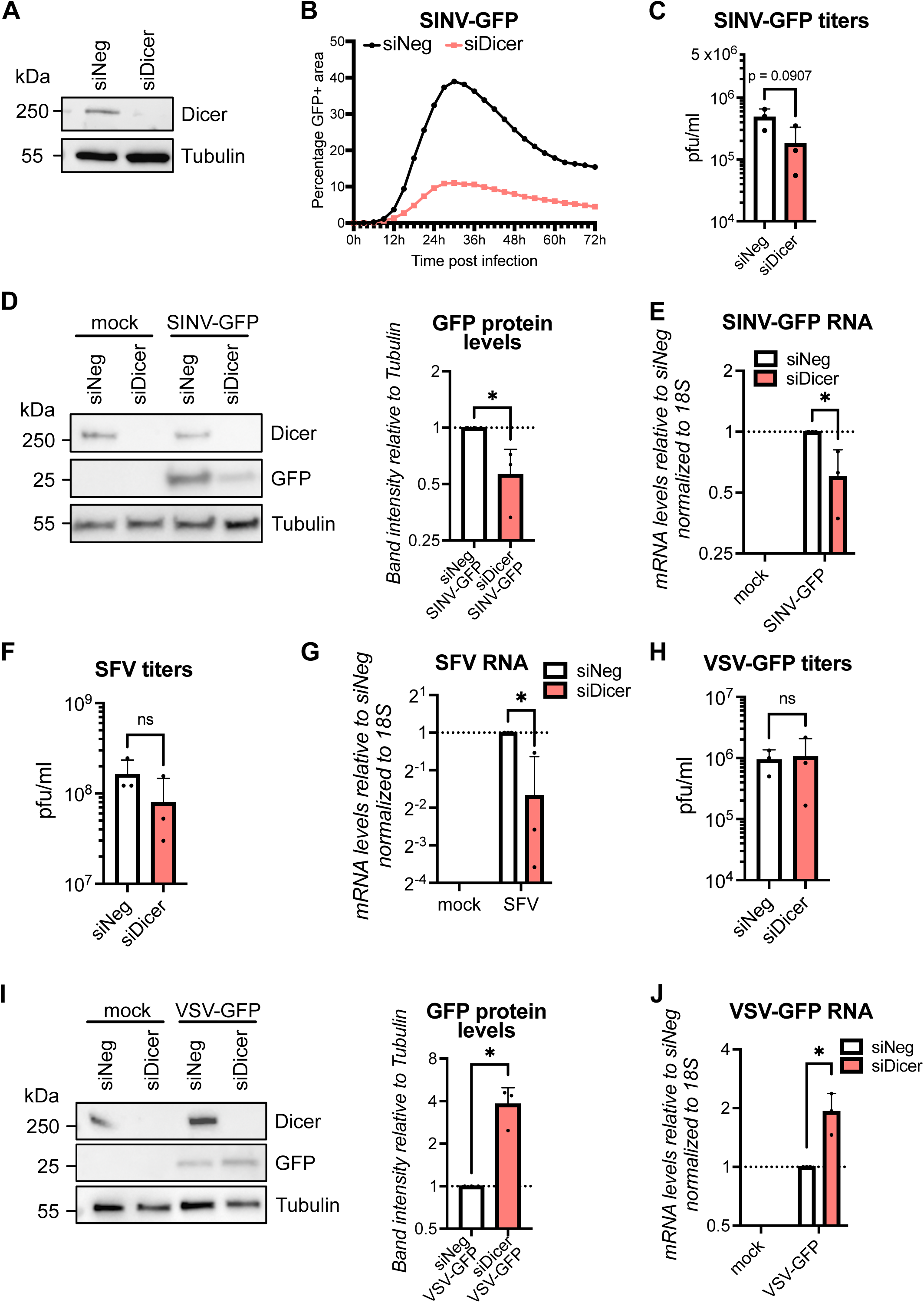
Dicer is proviral for SINV and SFV in *M. myotis* nasal epithelial cells. **(A)** Expression of Dicer was analyzed by western blot in *M. myotis* nasal epithelial cells after two rounds of siRNA targeting Dicer (siDicer) or a non-targeting control (siNeg). α-Tubulin was used as loading control. Images are representative of 2 independent experiments. **(B-J)** *M. myotis* nasal epithelial cells treated with siRNAs targeting Dicer (siDicer) or a non-targeting control (siNeg) were infected with SINV-GFP for 24h at MOI 0.2 **(B-E)**, SFV for 24h at MOI 20 **(F-G)** or VSV-GFP for 8h at MOI 1 **(H-J)**. **(B)** The number of GFP+ cells as a proxy for infection were monitored over time by live microscopy imaging. Percentages of GFP+ area were plotted for each condition as the average of two wells, six photos per well, for one experiment. **(C, F, H)** Supernatants were collected and viral titers were quantified by plaque assay. Means +/- standard deviation are plotted for 3 independent experiments. **(D, I)** Protein lysates were collected and analyzed by western blot using antibodies for GFP. α-Tubulin was used as loading control. n = 3 independent experiments. Quantification of GFP band intensities relative to Tubulin band intensities were quantified with ImageJ and the mean +/- standard deviation was plotted relative to siNeg mock samples for all three independent experiments. **(E, G, J)** RNAs were purified and levels of SINV RNA **(E)**, SFV RNA **(G)** or VSV RNA **(J)** were quantified by qRT-PCR. Means +/- standard deviation are plotted relative to siNeg infected samples for 3 independent experiments. For all graphs, p values were calculated using unpaired t test. * p<0.5 ; ns: not significant.

### *M. myotis* nasal epithelial cells only display a mild RNAi signature

Since *P. Alecto* and *T. brasiliensis* Dicer have enhanced antiviral RNAi activity, we wondered whether there was any evidence of antiviral RNAi activity in our *M. myotis* nasal epithelial cell line. We infected these cells with SINV-GFP and VSV-GFP, then sequenced small RNAs isolated from these cells. As control, we followed the same procedure with HEK293T cells, which we have previously showed display a weak RNAi signature against SINV-GFP (Girardi *et al*, 2013). Consistent with the lack of antiviral activity of mmDicer against SINV-GFP (**Fig. 1-2**), we did not observe a stronger RNAi signature against SINV-GFP in *M. myotis* cells than in HEK293T cells (**Fig. 3** and **S1**). This was also true for VSV-GFP (**Fig. 3** and **S1**), even though Dicer seemed to interfere with VSV-GFP at the RNA and protein level in *M. myotis* cells (**Fig. 2**). In both HEK293T and *M. myotis* cells, only a very low percentage of reads aligned to the SINV-GFP genome, and even lower to the VSV-GFP genome (**Fig. 3A**). Focusing on the SINV-GFP reads, while they showed an enrichment at 22nt on both strands in HEK293T cells, there was an enrichment of smaller size RNAs on the positive strand in the *M. myotis* cells, suggesting the presence of viral RNA degradation products (**Fig. 3B** and **S1A**). The same enrichment for smaller size RNAs on the positive strand could be seen for VSV-GFP reads in *M. myotis* cells, while almost no reads aligned to the VSV-GFP genome in HEK293T cells (**Fig. 3B** and **S1A**). The few viral reads that were 22nt long did display typical RNAi characteristics in both cell lines, as they aligned to both strands and throughout the viral genome (**Fig. 3C** and **S1B**). In both cell lines there was also an enrichment for paired positive and negative sense SINV-GFP reads that overlapped with a 2nt overhang, a signature of Dicer cleavage as part of the RNAi pathway (**Fig. 3D** and **S1C**), although that was not the case for VSV-GFP reads. However, the very low total number of viral reads of 22nt in both cell lines argues against a strong antiviral RNAi activity of mmDicer in *M. myotis* nasal epithelial cells against SINV and VSV. Altogether, our results indicate that mmDicer does not appear to have increased RNAi activity in *M. myotis* nasal epithelial cells.

**Figure 3:**
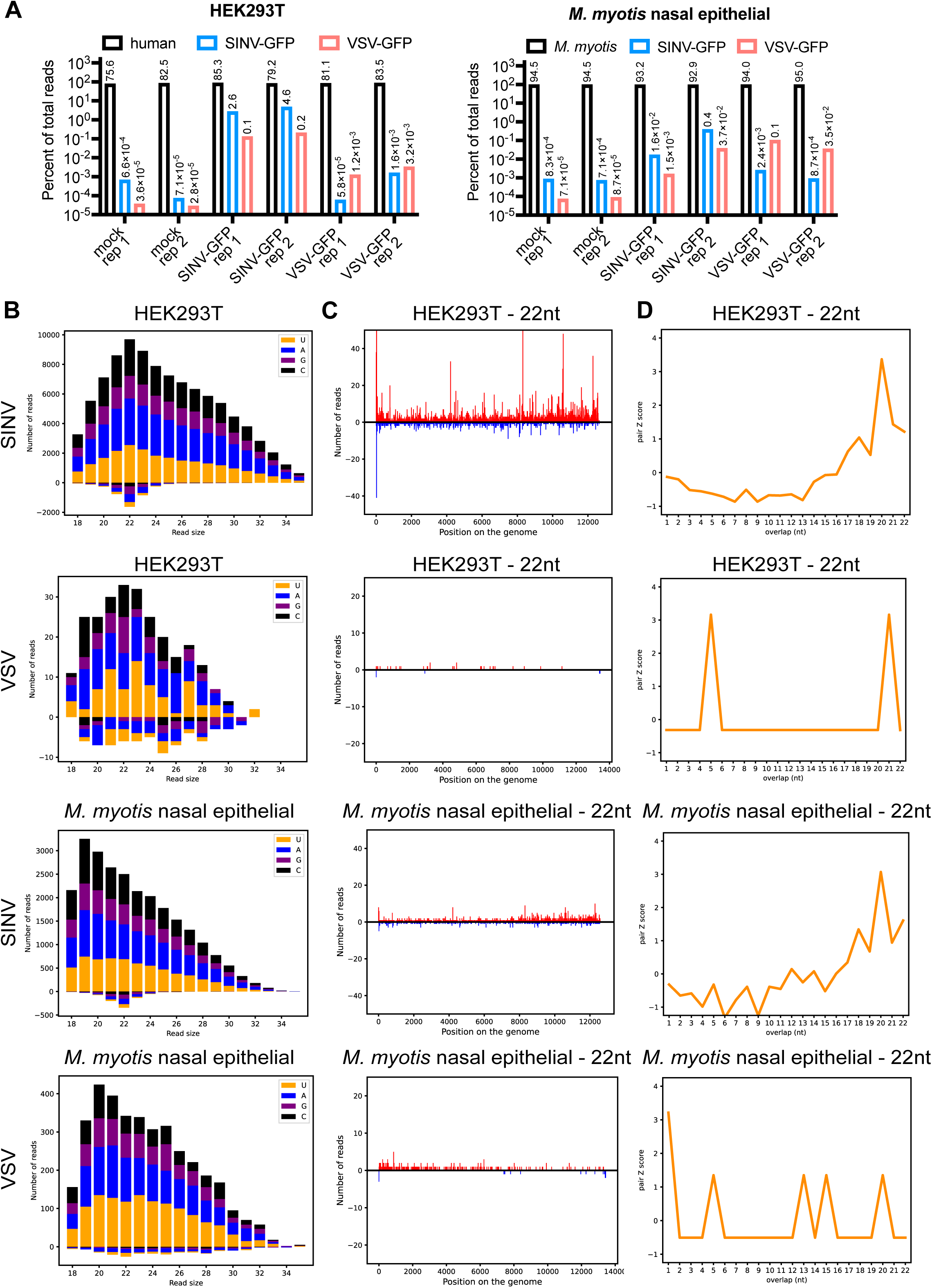
No strong RNAi signature in *M. myotis* nasal epithelial cells. HEK293T or *M. myotis* nasal epithelial cells were mock infected or infected with SINV-GFP or VSV-GFP, then small RNAs were extracted and sequenced. n = 2 independent experiments, see figure S1 for data from other replicate. **(A)** Percent of total sequencing reads that align to the human or *M. myotis* genome (black), the SINV-GFP genome (blue) or the VSV-GFP genome (red) for each sample in HEK293T cells (left) or *M. myotis* nasal epithelial cells (right). “rep” indicates the replicate number. **(B)** Number of reads that align to SINV-GFP or VSV-GFP for each cell type based on the read size. The total number of reads is further broken down into colors based on the identity of the first nucleotide of the read: yellow for U, blue for A, purple for G and black for C. **(C)** Location of the 22 nucleotide (nt) reads along the SINV-GFP or VSV-GFP genome for each sample. The number of reads falling on the same nucleotide is represented in red if the reads align to the genome (+ strand) and in blue if they align to the anti-genome (-strand). **(D)** For each sample, 22 nt small RNA pairs that overlap on + and - strands were analyzed and associated z-scores (Antoniewski, 2014) were plotted for the indicated nucleotide overlaps.

### Dicer relocalizes to cytoplasmic foci upon SINV infection in *M. myotis* nasal epithelial cells

To try to understand how mmDicer could play a proviral role in *M. myotis* nasal epithelial cells, we looked at the subcellular localization of Dicer upon SINV infection. In mock conditions, mmDicer exhibited a diffuse cytoplasmic localization, but interestingly upon WT SINV infection we saw a striking relocalization of Dicer into discrete cytoplasmic foci in *M. myotis* nasal epithelial cells. This relocalization could already be noticed at 8h (**Fig. 4A**) and was still apparent at 24h (**Fig. 4B**). There was no significant difference in both the number and size distribution of Dicer foci between 8h and 24h, although there was a small trend towards a decrease in the number of Dicer foci per cell (**Fig. 4C**). Importantly, we did not see any shift in Dicer localization upon SINV infection for HEK293T cells (**Fig. 4A-B**), as well as for the two other human cell lines we tested, human lung epithelial carcinoma A549 cells and human hepatocyte-derived carcinoma Huh7 cells (**Fig. S2A**). Overall, our results suggest a singular activity of Dicer during SINV infection in *M. myotis* nasal epithelial cells that leads to the relocalization of Dicer in large cytoplasmic foci, but that does not appear to occur in human cells.

**Figure 4:**
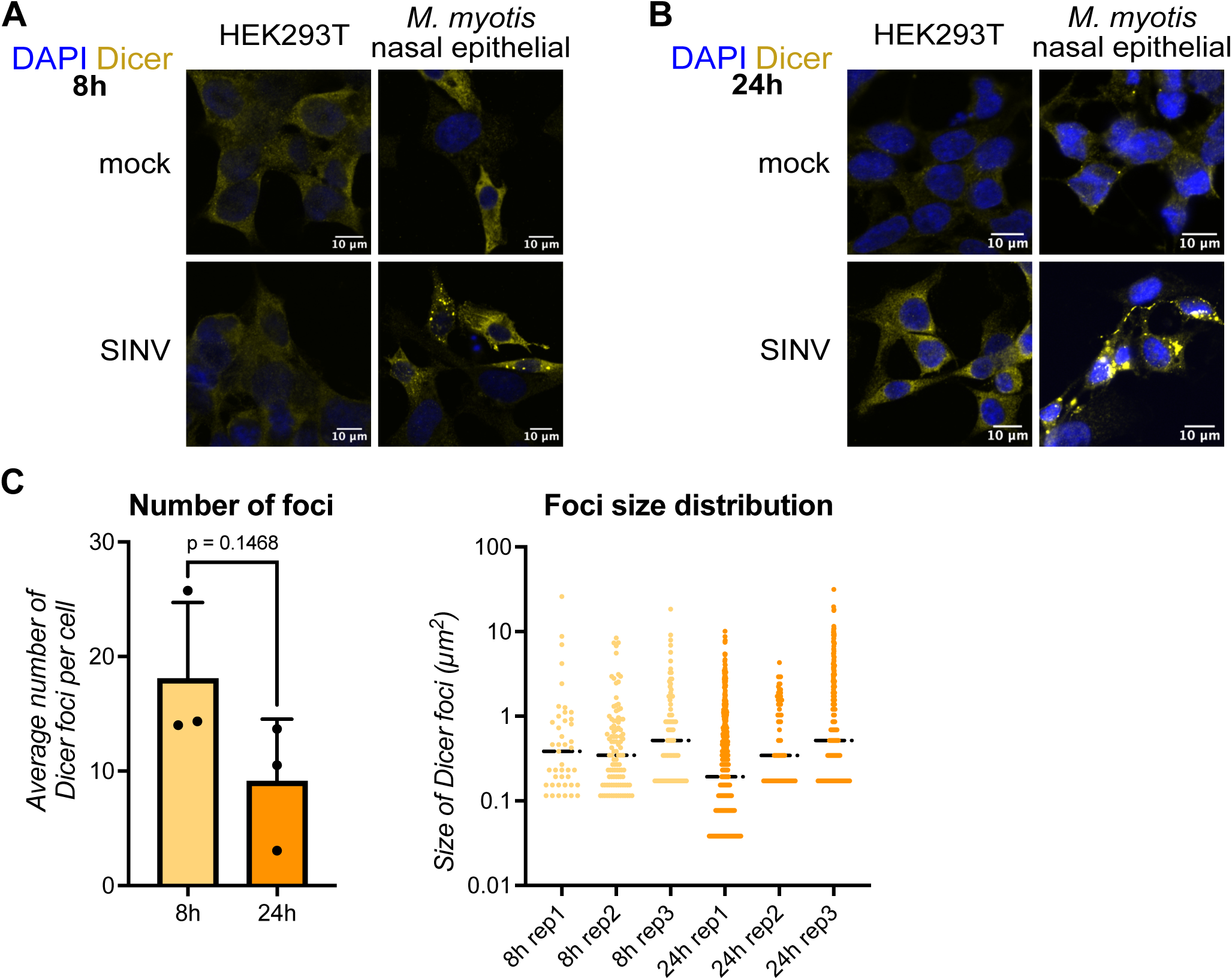
Dicer relocalizes to distinct cytoplasmic puncta upon infection in *M. myotis* nasal epithelial cells. **(A-C)** HEK293T were mock infected or infected with WT SINV for 8h at MOI 0.2 **(A)** or 24h at MOI 0.02 **(B)**. *M. myotis* nasal epithelial cells were mock infected or infected with WT SINV for 8h at MOI 5 **(A)** or 24h at MOI 2 **(B)**. Localization of Dicer (yellow) was imaged by immunofluorescence and confocal microscopy. DAPI staining (blue) indicates cell nuclei. Images are representative of at least 3 independent experiments. Scale bars 10µm. **(C)** For each condition, the number and size of Dicer foci in SINV infected *M. myotis* nasal epithelial cells were plotted for 3 or 4 independent experiments with each 1 or 2 fields containing approximately the same number of cells. The black horizontal bars represent the median. The p value was calculated using an unpaired t test.

### Dicer likely relocalizes to alphavirus viral factories in *M. myotis* nasal epithelial cells

SINV infected cells are known to accumulate large quantities of double stranded (ds)RNA in the cytoplasm as a result of replication (Griffin, 2007). Since dsRNA is the primary substrate of Dicer, we wondered if Dicer clustered around dsRNA in SINV infected cells. We infected *M. myotis* nasal epithelial cells with SINV and measured the co-localization of Dicer and dsRNA by confocal microscopy, using J2 antibodies that recognize dsRNAs longer than 40bp (Richardson *et al*, 2010). We saw that many of the Dicer foci co-localized with dsRNA in SINV infected cells (**Fig. 5A**), with a high Pearson correlation coefficient across multiple experiments (**Fig. 5B**), suggesting that Dicer forms foci in close proximity to dsRNA in *M. myotis* cells. As control, we could also see dsRNA accumulating in SINV infected HEK293T, A549 and Huh7 cells but Dicer staining remained diffuse and did not co-localize with dsRNA (**Fig. 5A-B, Fig. S2A**). Importantly, infection with the other alphavirus SFV also led to Dicer foci that co-localized with dsRNA (**Fig. 5A-B**). In contrast, infection with WT VSV is known to not accumulate dsRNA by J2 staining (Weber *et al*, 2006). And indeed, we did not detect dsRNA and there was no relocalization of Dicer upon infection in *M. myotis* cells (**Fig. S2B**). These results suggest that Dicer does not form large foci for every viral infection but its relocalization may be specific to alphaviruses or linked to dsRNA accessibility. dsRNA is often formed during viral replication, we thus wondered if Dicer was present at sites of SINV replication. Since J2 antibodies cannot distinguish between dsRNA of viral origin or originating from the host, we carried out fluorescence *in situ* hybridization (FISH) to specifically stain for SINV plus strand RNA. As expected, we saw discreet foci showing the accumulation of SINV RNA, likely at sites of viral replication, in both HEK293T cells and *M. myotis* nasal epithelial cells (**Fig. 5C**). Again, Dicer stayed diffuse upon infection in HEK293T cells. Importantly, the Dicer foci matched the SINV plus strand RNA foci in *M. myotis* cells (**Fig. 5C**). Altogether, our results argue for the recruitment of Dicer to sites of SINV viral replication.

**Figure 5:**
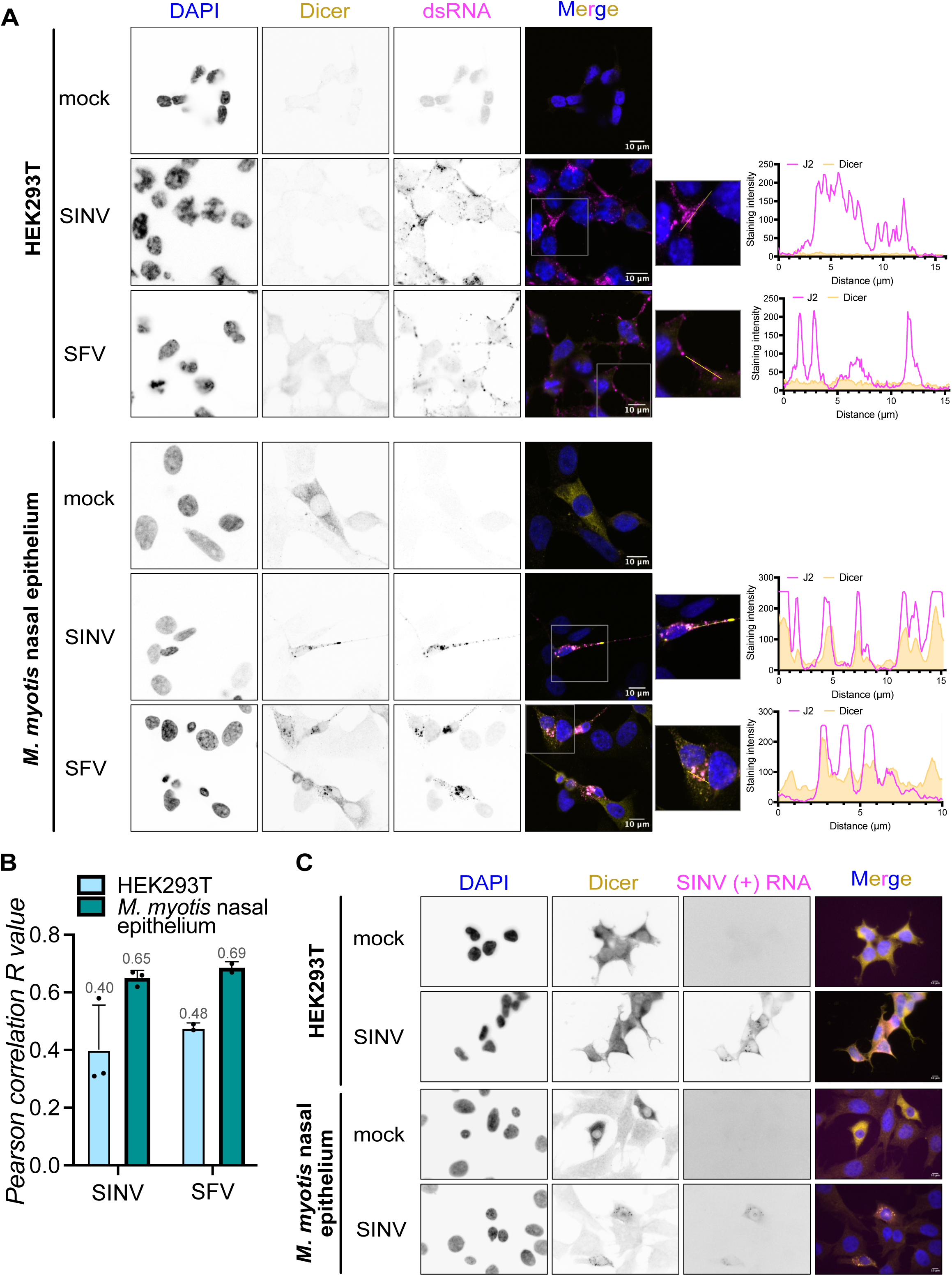
Dicer likely relocalizes to SINV viral factories in *M. myotis* nasal epithelial cells. **(A-B)** HEK293T were mock infected or infected with WT SINV or SFV for 24h at MOI 0.02. *M. myotis* nasal epithelial cells were mock infected or infected with WT SINV or SFV for 24h at MOI 2 or 5 respectively. Localization of Dicer (yellow) and dsRNA via J2 antibodies (magenta) was imaged by immunofluorescence and confocal microscopy. DAPI staining (blue) indicates cell nuclei. Images are representative of 3 (SINV) or 2 (SFV) independent experiments. Scale bars 10µm. Co-localization between Dicer and dsRNA signals was quantified and Pearson correlation R coefficients were calculated and plotted for 3 (SINV) or 2 (SFV) independent experiments – mean (numerical value indicated in gray font above each bar) +/- standard deviation **(B)**. A zoomed-in image of the boxed area in some merge images was added. The graphs show the signals for each channel along the line drawn in the zoomed in images. **(C)** HEK293T or *M. myotis* nasal epithelial cells were mock infected or infected with WT SINV for 24h at MOI 0.02 or 4, respectively. Localization of Dicer (yellow) by immunofluorescence and of SINV plus strand (+) RNA (magenta) by FISH was imaged with a fluorescent microscope. DAPI staining (blue) indicates cell nuclei. Images are representative of 2 independent experiments. Scale bars 10µm.

### *M. myotis* Dicer alone is not sufficient for relocalization in NoDice cells

Finally, we wondered if there was something inherently special about mmDicer that triggered its relocalization upon SINV infection. To test this, we went back to our human NoDice cells expressing mmDicer (**Fig. 1**). Immunoprecipitating mmDicer in these cells showed that mmDicer still interacted with known hDicer interactors such as PACT and TRBP in mock conditions, as well as PKR and ADAR1, which we previously showed interact with hDicer during SINV infection (Montavon *et al*, 2021) (**Fig. 6A**). As expected, while J2 staining showed discrete foci characteristic of SINV infection, hDicer stayed diffuse throughout the cytoplasm (**Fig. 6B**). Importantly, there was also no relocalization of mmDicer upon SINV infection (**Fig. 6B**). These results suggest that mmDicer alone is not sufficient to form large foci, and that factors specific to *M. myotis* nasal epithelial cells are required for relocalization.

**Figure 6:**
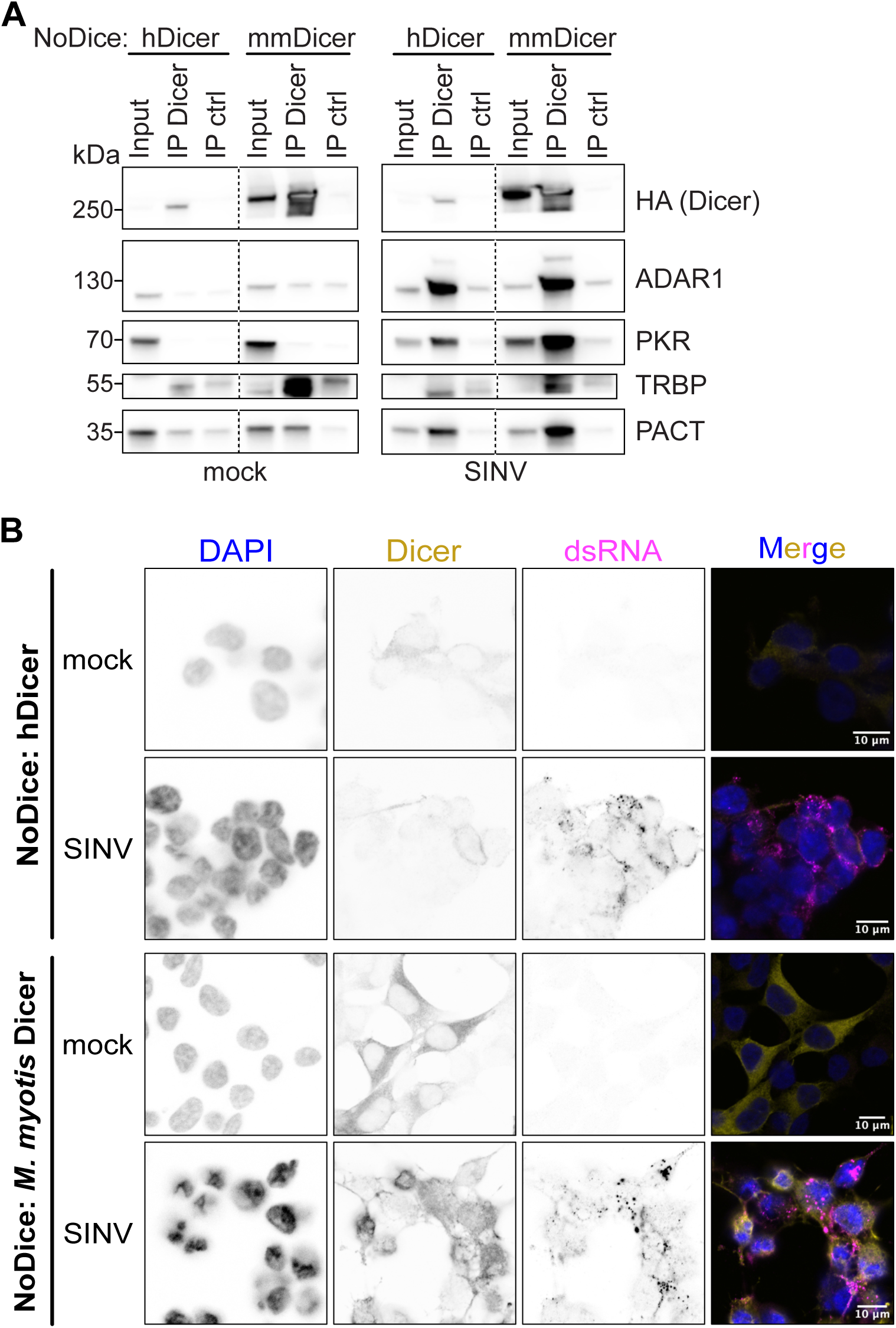
*M. myotis* Dicer alone is not sufficient for relocalization. HEK293T NoDice cells expressing HA-tagged hDicer or mmDicer were mock infected or infected with SINV-GFP for 24h at MOI 0.02. **(A)** Cells were then lysed, a small aliquot was kept as input, and Dicer was immunoprecipitated with anti-HA coated magnetic beads (IP Dicer) or anti-myc coated beads (IP ctrl). Eluted proteins were analyzed by western blot using antibodies for the known hDicer interactors ADAR1, PKR, TRBP and PACT. Anti-HA antibodies show efficient immunoprecipitation of hDicer and mmDicer. Images representative of 2 independent experiments. **(B)** Localization of Dicer (yellow) and dsRNA via J2 antibodies (magenta) was imaged by immunofluorescence and confocal microscopy. DAPI staining (blue) indicates cell nuclei. Images are representative of 2 independent experiments. Scale bars 10µm.

## Discussion

Recent studies have highlighted a role for Dicer in bat antiviral immunity and tolerance to viruses through RNAi in two different bat species, *P. alecto* and *T. brasiliensis* (Dai *et al*, 2024; Owolabi *et al*, 2025). In this study, we set out to characterize the antiviral activity of Dicer in *M. myotis* nasal epithelial cells. Interestingly, mmDicer did not protect from SINV and VSV infection when expressed in a human Dicer knock out cell line (**Fig. 1**). Moreover, knocking down Dicer in *M. myotis* nasal epithelial cells led to a decrease in SINV and SFV infection, while results were more mitigated with VSV (**Fig. 2**). Consistent with these results, we did not find a strong antiviral RNAi signature when analyzing the viral small RNA profiles of SINV and VSV infected *M. myotis* nasal epithelial cells (**Fig. 3** and **S1**). However, we did observe a striking relocalization of Dicer in large cytoplasmic foci upon SINV and SFV infection in *M. myotis* nasal epithelial cells that we did not witness in human cells (**Fig. 4** and **S2**). These Dicer foci are in close proximity with dsRNA and SINV plus strand RNA, suggesting that Dicer is recruited to sites of SINV replication (**Fig. 5**). Finally, mmDicer does not form these foci when expressed in human Dicer knock out cells, suggesting that its relocalization requires specific factors that are present in *M. myotis* nasal epithelial cells (**Fig. 6**).

Our results that mmDicer does not have increased antiviral RNAi activity is unexpected in view of the recently published results on *P. alecto* and *T. brasiliensis* Dicer, especially as SINV was also used in the *P. alecto* study (Dai *et al*, 2024; Owolabi *et al*, 2025). This discrepancy could come from differences in bat species, as *P. alecto* and *M. myotis* are thought to have diverged more than 60 million years ago, and *T. brasiliensis* and *M. myotis* at least 35 million years ago (Teeling *et al*, 2005; Agnarsson *et al*, 2011; Hao *et al*, 2024). Since in human and mouse cells the antiviral RNAi activity of Dicer can be cell-type dependent, it is also possible that the activity of bat Dicer is influenced by the cell type used in each study, especially if different bat species and cell types have differences in IFN activity (Geng *et al*, 2024; Bondet *et al*, 2021; Clayton & Munir, 2020). In mammalian cells, the same Dicer protein could play a role in antiviral immunity through its RNAi activity, but also through direct regulation of the innate immune reponse and through the maturation of miRNAs that regulate the innate immune response (Gaucherand *et al*, 2024; Maillard *et al*, 2019). These multiple interconnected activities of Dicer make the study of Dicer antiviral activity difficult. We employed a transient siRNA knockdown strategy instead of knocking out Dicer to limit the impact of removing Dicer on the expression of miRNAs, which have a notoriously long half-life (Marzi *et al*, 2016), but there could still be indirect effects of Dicer influencing its propensity to be anti- or proviral in specific cellular contexts. In general, it will be important to study the activity of Dicer in more bat species to resolve this discrepancy.

The case of VSV is interesting, as knocking down Dicer in *M. myotis* cells led to an increase in VSV-GFP RNA and proteins levels (**Fig. 2F-G**). Yet, there was almost no viral siRNAs (**Fig. 3, Fig S1**). The mechanism behind this apparent antiviral activity of Dicer is thus unclear. Nonetheless, it serves as a good control of a virus that does not trigger relocalization of Dicer in *M. myotis* cells, suggesting that this relocalization is context-specific and not an artefact of infection. Since dsRNA is not detectable upon VSV infection unlike infection with alphaviruses, it is tempting to hypothesize that the relocalization of Dicer is linked to dsRNA accessibility. It will be interesting to test this hypothesis.

Dicer relocalization to discreet cytoplasmic foci upon infection in *M. myotis* cells is reminiscent of *Drosophila* D2 bodies (Nishida *et al*, 2013). These D2 bodies are cytoplasmic foci containing Dicer-2 and its co-factor R2D2 that are important for the processing of endogenous siRNAs in *Drosophila* cells. Yet, our small RNAseq results did not identify such activity of Dicer. Moreover, we see a proviral effect instead of an antiviral effect associated with Dicer relocalization, although it remains purely correlative as we do not have concrete evidence so far that the two are directly linked. We can nonetheless propose several hypotheses as to how the relocalization of Dicer could play a proviral role. Potential consequences of this relocalization could be the sequestration of Dicer by SINV, for example to prevent Dicer activity as part of the antiviral RNAi pathway. Alternatively, Dicer could be recruited to sites of SINV replication to hide dsRNA from PRRs or to directly tone down the local innate immune response. One possible explanation for this recruitment, which remains to be tested, could be that mmDicer has a higher affinity for dsRNA than its human counterpart in *M. myotis* cells, maybe thanks to specific co-factors, but with no increased cleavage activity. Dicer could also have a structural role to help viral factory formation and alphavirus replication. More work will be needed to identify the purpose of Dicer relocalization and the link between this relocalization and Dicer proviral activity.

It is intriguing that this relocalization does not appear to occur in human cells, or at least in the human cell lines that we have tested. The fact that mmDicer alone is not sufficient for relocalization is consistent with mmDicer and hDicer being very similar, with 95% conservation between the two proteins at the amino acid level. Work to identify the *M. myotis* cell specific factors responsible for Dicer relocalization is ongoing. Host proteins that only interact with viral proteins in bat cells are good candidates. On top of finding the mechanism of Dicer relocalization, it will also be important to determine whether Dicer forms cytoplasmic foci during infection with other viruses and in other stress conditions. It will especially be important to test viral strains that are more biologically relevant to *M. myotis*. While many different alphaviruses have been detected in several species of bats in the wild, including SINV and SFV, it is still not clear whether bats constitute a reservoir for alphaviruses as this has not been well studied (Fagre & Kading, 2019). Finally, it would be interesting to determine if Dicer relocalization occurs in other bat species and cell types, and whether this relocalization could explain the discrepancy in the observed antiviral RNAi activity of Dicer.

## Materials and methods

### Cell culture and viral infections

SV40 transformed *M. myotis* nasal epithelial cells (He *et al*, 2014b) were a kind gift from Bernd Köllner (FLI, Greifswald, Germany) through Lucie Etienne (CIRI, Lyon, France). HEK293T/NoDice (2.20) cells (Bogerd *et al*, 2014) were a kind gift from Bryan Cullen (Duke University, Durham NC, USA). All cells were maintened in Dulbecco’s modified Eagle medium (DMEM, Gibco, Life Technologies) supplemented with 10% fetal bovine serum (FBS, bioSera FB1090/500, Batch# S00O0) in a humidified atmosphere of 5% CO_2_ at 37°C.

WT SINV (strain AR339) and SINV-GFP (kindly provided by M. C. Saleh, Institut Pasteur, Paris, France, Saleh et al., 2009) viral stocks were produced and amplified in BHK21 cells. SFV (strain UVE/SFV/UNK/XX1745; EVAg 001V-02468) was propagated in Vero E6 cells from the initial stock. WT VSV (strain Indiana isolate PI10) and VSV-GFP (Mueller *et al*, 2010) were propagated in BHK21 cells from the initial stock. Stock titrations were performed by plaque assay on VERO E6 cells. Cells were infected at MOIs ranging from 0.02 to 20, for 8h or 24h, as indicated in the figure legends.

### Plasmids and cloning

pLenti6 Flag-HA-V5 vector was modified from pLenti6-V5 gateway vector (Thermo Fisher scientific V49610) by Gibson cloning (New England Biolabs) as previously described (Baldaccini *et al*, 2024), then further modified to remove the V5 tag and ccdB cassette. plenti6 Flag-HA hDicer was already described (Baldaccini *et al*, 2024). mmDicer was cloned from *M. myotis* nasal epithelial cells mRNA using Superscript IV one-step RT-PCR (Thermo Fisher Scientific) with primers Forward (CAGTGTGGTGGAATTCTGCAGAAAA GCCCTGCTTTGCAACCC) and Reverse (CCAAACTCATTACTAACCGGTTCAGCTGCT GGGAACCTGAG). plenti6 Flag-HA-V5 was digested with AgeI and PstI restriction enzymes, then ligated with amplified mmDicer with matching overhangs by Gibson cloning (New England Biolabs). Catalytic mutations (E1562A and E1819A) were introduced by PCR inside primers Forward (GACTGCGTGGCTGCCCTGCTGGGCTGC TATTTAA) + Reverse (AGCAGGGCAGCCACGCAGTCGGCGATGCTTTTGT) and Forward (GGGATATTTTTGCTTCACTTGCTGGTGCCATTTAC) + Reverse (GCAAGTGA AGCAAAAATATCCCCCATGGCCTTG), then ligated to AgeI and PstI digested plenti6 Flag-HA-V5 by Gibson cloning (New England Biolabs).

### Lentivirus production and generation of stable cell lines

HEK23T cells were plated in 6 well plates and transfected with 0.33 µg pVSV envelope plasmid (Addgene #8454) and 1.33 µg pSPAX2 packaging plasmid (Addgene #12260), and 1.7 µg of either plenti6 Flag-HA Empty, plenti6 Flag-HA hDicer, plenti6 Flag-HA mmDicer or plenti6 Flag-HA mmDicer CM, using Lipofectamine 2000 reagent (Invitrogen, Fisher Scientific) according to the manufacturer’s protocol. Media was replaced 6h later, then the supernatent containing viral particles was collected at 48h and filtered through a 0.45 µm PES filter. HEK293T NoDice (Bogerd *et al*, 2014) were then transduced with the filtered lentiviruses mixed with 4µg/ml polybrene. Efficiently transduced cells were selected for several days with blasticidine at 15 µg/mL and polyclonal populations were used for experiments.

### Dicer knock down

*M. myotis* Dicer specific siRNAs were designed by siTOOLs Biotech to form a siPOOL of 30 siRNAs. A siPOOL of 30 non-targeting siRNAs was used as control (siTOOLs Biotech). *M. myotis* nasal epithelial cells were reverse transfected with siNeg or siDicer at 3nM final siRNA concentration using Lipofectamine RNAiMax (ThermoFisher) following siTOOLs Biotech instructions. Cells were transfected again the following day with siNeg or siDicer at 3nM final siRNA concentration. The next day, cells were collected to check for efficient knock down or infected with different viruses.

### Live cell imaging

*M. myotis* nasal epithelial cells were treated with two rounds of siRNAs then infected with SINV-GFP at MOI 0.2. Cells were then placed into a CellCyte X live cell imaging system (Cytena) where GFP fluorescence and phase contrast pictures (10X objective) of every well were taken every 3h for 72h. The percentage of GFP+ area over the total cell area was quantified with the CellCyte Studio software and averaged from six photos of 2 individual wells per condition.

### Protein extraction and western blot

Proteins were harvested and homogenized in appropriate quantity of ice cold lysis buffer (50 mM Tris-HCl pH 7.5, 150 mM NaCl, 5 mM EDTA, 1% Triton X-100, 0.5% SDS protease free inhibitor tablet (complete Mini; Sigma Aldrich)). Protein amounts were quantified using Bradford reagent (Bio-Rad), diluted in lysis buffer + Laemmli and boiled for 5 min before loading. Proteins were loaded on 10% acrylamide-bis-acrylamide gels, or 4–20% Mini-PROTEAN TGX Precast Gels (Bio-Rad) for the Dicer immunoprecipitation experiment. Proteins were separated by migration at 140 V in 1X Tris-Glycine-SDS buffer (Euromedex), then electro-transferred on a nitrocellulose membrane (Amersham 0.45 µm) in 1X Tris-Glycine buffer supplemented with 10% ethanol. Appropriate loading and transfer efficiency were verified by Ponceau S staining (Merck). Membranes were blocked in 5% milk diluted in PBS-Tween 0.2%. Membranes were stained with primary antibodies overnight at 4 °C at the following dilutions: anti-DICER (1:2000, A301-937A, Euromedex, Bethyl), anti-HA-HRP (1:1000, 12013819001 Merck, Sigma Aldrich), anti-GFP (1:1000, 11814460001 Merck, Sigma Aldrich), anti-SINV nsp2 (1:5000, kind gift from Diane Griffin, Johns Hopkins University School of Medicine, Baltimore, MD), anti-PKR (1:1000, ab32506 Abcam), anti-PACT (1:500, ab75749 Abcam), anti-TRBP (1:500, sc-514124, Cliniscience, Santa Cruz), anti-ADAR1 (1:1000, E6X9R #81284 Cell Signaling), anti-α-Tubulin-HRP (1:10000, Sigma, T6557). After several washes with PBS-Tween 0.2%, the membranes were stained with the following secondary antibodies coupled to horseradish peroxidase (HRP): anti-mouse-HRP (1:4000, A4416 Merck, Sigma-Aldrich) or anti-rabbit-HRP (1:10,000, NA9340V Cytiva, Sigma-Aldrich). Detection was carried out using SuperSignal West Femto maximum sensitivity substrate (Pierce, Fisher Scientific) and visualized with a Fusion FX imaging system (Vilber).

### Plaque assay

For SINV-GFP infections, Vero E6 cells were seeded in 96-well plates and infected with 10-fold serial dilutions of infection supernatants. After 1 hour, the inoculum was removed and cells were covered with 2.5% carboxymethyl cellulose and cultured for 72 hours at 37°C in a humidified atmosphere of 5% CO2. Plaques were counted manually under the microscope. For SFV and VSV-GFP infections, Vero E6 cells were seeded in 12-well plates and infected with 10-fold serial dilutions of infection supernatants. After 1 hour, the inoculum was removed, cells were covered with 1.2% Avicel overlay (vivapur mcg, JRS Pharma) and cultured for 48 hours at 37°C in a humidified atmosphere with 5% CO2. Cells were then washed and fixed with 4% formaldehyde (Merck, Sigma Aldrich) in PBS (phosphate buffered saline, Gibco) for 20 minutes, then stained with crystal violet for 5 minutes (Sigma-Aldrich). Plates were washed with water and plaques were counted manually. Viral titers were calculated according to the formula: PFU/mL = #plaques/ (Dilution*Volume of inoculum).

### RNA and small RNA extractions

Total RNA was extracted from cell lysates using Trizol Reagent (15586018, InVitrogen) to which 200 µl of chloroform was added before centrifugation for 15 minutes at 12,000 xg at 4°C. For regular RNA extraction, the aqueous layer was collected and nucleic acids were precipitated in an equal volume of isopropanol for 10 min at room temperature. For small RNA extraction, the aqueous layer was collected and nucleic acids were precipitated in 3 volumes of 100% ethanol + 10% 1M sodium acetate for 30 minutes at -80°C. In both cases, precipitated RNA was then pelleted for 10 min at 12,000 xg at 4°C then washed with 70% ethanol and reconstituted in RNase free water.

### Northern blot analysis

3 µg of total RNA was loaded onto a 17% urea acrylamide gel and electro separated in a 1X Tris-Borate-EDTA solution. The RNA was then electro transferred to a Hybond-NX membrane (GE Healthcare), and chemically cross linked to the membrane with 1-ethyl-3-[3-dimethylaminopropyl]carbodiimide hydrochloride (EDC) (Sigma Aldrich) for 90 min at 65°C. Membranes were then pre-hybridized for 30 min with PerfectHyb plus (Sigma Aldrich) at 50°C. Probes consisting of an oligodeoxyribonucleotide sequence for which the 5′-was labelled with 25 μCi of [γ-32P]dATP using T4 polynucleotide kinase (Thermo Fisher Scientific) were incubated overnight with the membranes at 50°C. The membranes were then washed twice at 50°C for 10 min with 5X SSC/0.1% SDS, then for 5 min with 1X SSC/0.1% SDS. Finally, membranes were placed in contact with a phosphorimager plate overnight and imaged with a Bioimager FLA-7000 scanner (Fuji). Probe sequences: miR-16 (CGCCAATATTTACGTGCTGCTA), U6 (GCAGGGGCCATGCTAATCTTCTCTGT ATCG).

### RT-qPCR analysis

After RNA extraction, samples were incubated with DNase I (Invitrogen, Fisher Scientific) for 20 min at 37°C. The RNA was then reverse-transcribed into cDNA using SuperScript IV (SSIV) reverse transcriptase (Invitrogen, Fisher Scientific) or AMV reverse transcriptase (Promega) with a random nonameric primer, following the manufacturer’s protocol. Quantitative real-time PCR was performed using SYBR Green PCR Master Mix (Fisher Scientific) and the following primers: 18S F (CTCTTAGCTGAGTGTCCCGC), 18S R (CTTAATCATGGCCTCAGTTCCGA), SINV F (CCACTACGCAAGCAGAGACG), SINV R (AGTGCCCAGGGCCTGTGTCCG), SFV F (GCGGCAAGAAGGAGAACTGC), SFV R (CAAGCGAAAGCCTCGTCCAC), VSV F (TGTACAATCAGGCGAGAGTC), VSV R (GGAACTCATGATGAATGGATTGGG).

### Small RNA sequencing and bioinformatic analysis

After small RNA extraction, samples were ran on a Bioanalyzer chip to confirm RNA integrity. Samples were then shipped to BGI for library preparation with unique molecular identifiers, and sequenced as single-end 50 base reads on their DNBSEQ sequencing technology platform with at least 20 million reads per sample.

Raw reads were pre-cleaned by the sequencing platform, including adapter, contaminant and low-quality removal (Phred+33). Before alignment, redundant sequences were collapsed to one copy using fastq_collapser from FASTX-Toolkit 0.0.14 (http://hannonlab.cshl.edu/fastx_toolkit/) and a read-size filter was applied (18-35 bp). HEK293T and *M. myotis* short reads were then aligned to the human genome assembly GRCh38 (GCF_000001405.40) and the *M. myotis* genome mMyoMyo1.p (GCF_014108235.1), respectively. Mapping was performed using Bowtie 1.3.1 (Langmead *et al*, 2009) with parameters ‘-v 1 -a --best --strata’. Unmapped reads were next separately aligned against virus genomes SINV-GFP (private sequence derived from NC_001547.1 – RefSeq database) and VSV-GFP (FJ478454.1) using the following parameters ‘-v 2 -y -q -m 30 -a --best --strata’. Finally, siRNA signature was examined on 22nt mapped reads by the python tool signature by (Antoniewski, 2014) with options ‘22 22 1 22’. Sequencing data have been deposited on NCBI’s Gene Expression Omnibus (Edgar *et al*, 2002) and are accessible through GEO Series accession number GSE301457.

### Immunostaining

Cells were seeded onto Labtek eight-chamber slides (Nunc Lab Tek Chamber Slide System) at a density of 8,000 cells per well. The next day, cells were infected for 8h or 24h with WT SINV, WT VSV or SFV. Cells were then fixed for 10 minutes at room temperature with 4% formaldehyde (Sigma-Aldrich) diluted in 1X PBS. Blocking was carried out with 5% normal goat serum in 0.1% Triton X-100 in 1X PBS for 1 hour at room temperature. Primary antibody staining was carried out for 3h at room temperature using the following antibodies diluted in blocking buffer: anti-Dicer (1:1000, A301-937A, polyclonal, Bethyl) and J2 anti-dsRNA (1:1000, RNT-SCI-10010200, Jena Bioscience). The cells were then incubated for 1 hour at room temperature with secondary antibodies goat anti-mouse Alexa 594 (A11032, Invitrogen) or goat anti-rabbit Alexa 488 (A11008, Invitrogen), both diluted at 1:1000 in 1X PBS with 0.1% Triton X-100. Between each step, cells were washed thoroughly with 1X PBS containing 0.1% Triton X-100. Nuclei were stained with DAPI (4’,6-diamidino-2-phenylindole, dichlorhydrate, D1306, Thermo Fisher) at 1:5000 in 1X PBS for 5 minutes. Finally, the slides were mounted with coverslips using Fluoromount-G mounting media (Southern Biotech) and observed under a confocal microscope (LSM780, Zeiss).

### Sequential immunostaining and FISH

Mock- or WT SINV-infected cells were cultured on 18 mm round coverslips placed in 12-well plates. Cells were fixed with 3,7% formaldehyde (Sigma-Aldrich) diluted in 1X PBS for 10 minutes at room temperature. Following fixation, cells were permeabilized with 0.1% Triton X-100 in 1X PBS for 5 minutes at room temperature. Immunostaining was performed as described in the previous section but with 1h incubation steps for primary and secondary antibody staining. Subsequently, cells were fixed again with 3.7% formaldehyde (Sigma-Aldrich) diluted in 1X PBS for 10 minutes at room temperature. For RNA detection, cells were incubated overnight at room temperature with SINV genome-specific Stellaris Cal Fluor Red 610 RNA FISH Probes (LGC Biosearch Technologies, custom made by Erika Girardi (Messmer *et al*, 2024)), kindly provided by Erika Girardi (IBMC, Strasbourg, France). Probes were diluted in RNA FISH hybridization buffer (Stellaris, Biosearch Technologies). Nuclei were stained with DAPI (D1306, Invitrogen, Thermo Fisher Scientific) for 30 minutes. Finally, coverslips were mounted using Fluoromount-G mounting medium (Invitrogen, Thermo Fisher Scientific), and the samples were imaged by fluorescence microscopy (Olympus BX51).

### Image analysis

All image analyses were performed on confocal images and using the Fiji software (Schindelin *et al*, 2012). Detection thresholds and all parameters were standardized between the different images and cell lines to ensure consistency in the analysis. Each analysis was performed on two (SFV) or three (SINV) biological replicates, with each field containing approximately the same number of cells. The “Plot profile” tool on Fiji was used to visualize fluorescent intensity as a function of distance for each channel. The selection of the line or field of observation was made arbitrarily to best represent the whole slide. Threshold parameters were adjusted accordingly to measure the size of various objects, and the minimum size accepted was set to 0.1 μm² to limit background detection. The “Analyse particle” tool was used to extract numerical data, which was then systematically analyzed. The colocalization analysis between two fluorescent signals was performed using the "Colocalization Finder" tool in Fiji. This tool analyzes colocalization on a pixel-by-pixel basis by comparing the intensity of each pixel in one channel to the corresponding pixel in the second channel of a bi-color image. Importantly, the thresholds or brightness levels in the image are not changed during this process. The analysis generates a scatter plot, from which a Pearson correlation coefficient is calculated. A value of +1 indicates perfect correlation, 0 indicates no correlation, and -1 signifies perfect anti-correlation.

### Dicer immunoprecipitation

NoDice cells expressing different HA-tagged Dicer constructs were plated and mock-infected or infected with SINV-GFP at MOI 0.02. 24 hours later, cells were harvested, washed twice with ice-cold 1X PBS (Gibco, Life Technologies), and resuspended in lysis buffer (50 mM Tris-HCl pH 7.5, 140 mM NaCl, 6 mM MgCl2, 0.1% NP-40), supplemented with a Complete-EDTA-free Protease Inhibitor Cocktail tablet (complete Mini; Sigma Aldrich). Cells were lysed on ice for 10 min and debris were removed by 10 min centrifugation at 12,000 *xg* at 4°C. An aliquot of the cleared lysates was kept aside as protein input. The rest of the samples were divided in two equal volumes and incubated with 40 μL magnetic microparticles coated with monoclonal anti-HA or anti-MYC antibodies (MACS purification system, Miltenyi Biotech) at 4°C for 1 hour with constant rotation. Next, samples were passed through μ Columns (MACS purification system, Miltenyi Biotech) and washed four times with 200 μL lysis buffer. Finally, proteins were eluted with 2X Laemmli buffer (10% glycerol, 4% SDS, 62.5 mM TrisHCl pH 6.8, 5% (v/v) 2-β-mercaptoethanol, Bromophenol Blue) that was pre-warmed to 95 C°.

## Acknowledgements

We would like to thank members of the laboratory for insightful discussions and suggestions. This work of the Interdisciplinary Thematic Institute IMCbio+, as part of the ITI 2021-2028 program of the University of Strasbourg, CNRS and Inserm, was supported by IdEx Unistra (ANR-10-IDEX-0002), by SFRI-STRAT’US project (ANR-20-SFRI-0012), EUR IMCBio (IMCBio ANR-17-EURE-0023) under the framework of the French Investments for the Future Program. It was also supported by an Agence Nationale pour la Recherche PRCI grant to SP (ANR-21-CE35-0018-01). LG was funded by a postdoctoral fellowship from the Fondation pour la Recherche Médicale (funding number SPF202209015746).

---

**Figure S1:**
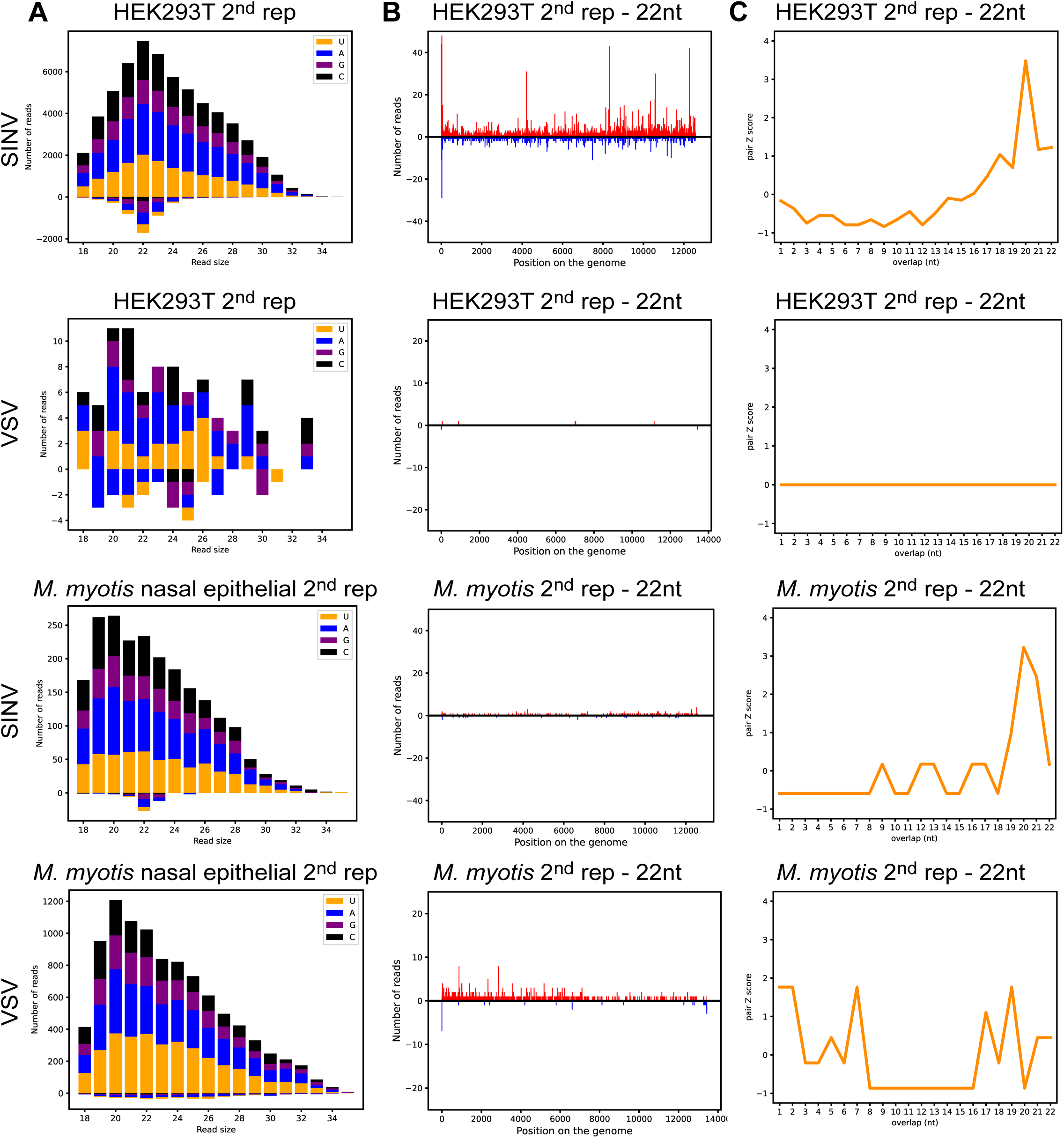
Small RNAseq results for the rest of HEK293T and *M. myotis* nasal epithelial samples. HEK293T or *M. myotis* nasal epithelial cells were mock infected or infected with SINV-GFP or VSV-GFP, then small RNAs were extracted and sequenced. n = 2 independent experiments, see figure 3 for data from other replicate. **(A)** Number of reads that align to SINV-GFP or VSV-GFP for each cell type based on the read size. The total number of reads is further broken down into colors based on the identity of the first nucleotide of the read: yellow for U, blue for A, purple for G and black for C. **(B)** Location of the 22 nucleotide (nt) reads along the SINV-GFP or VSV-GFP genome for each sample. The number of reads falling on the same nucleotide is represented in red if the reads align to the genome (+ strand) and in blue if they align to the anti-genome (-strand). **(C)** For each sample, 22 nt small RNA pairs that overlap on + and - strands were analyzed and associated z-scores (Antoniewski, 2014) were plotted for the indicated nucleotide overlaps.

**Figure S2:**
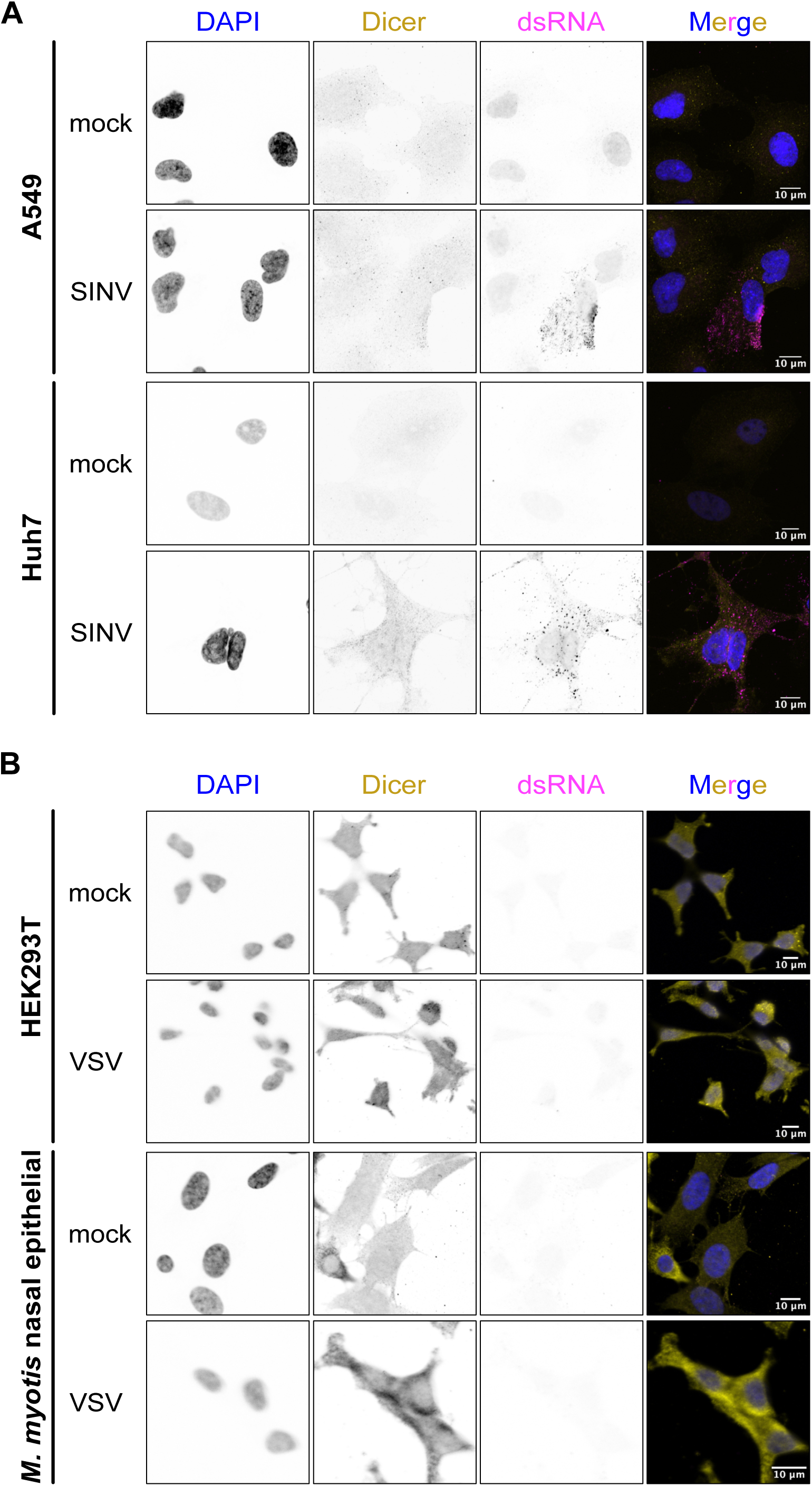
No Dicer foci are found in other human cell lines and in VSV infected cells. **(A)** Human A549 and Huh7 cells were mock infected or infected with WT SINV at 24h at MOI 0.2 and 0.02, respectively. Localization of Dicer (yellow) and dsRNA via J2 antibodies (magenta) were imaged by immunofluorescence and confocal microscopy. DAPI staining (blue) indicates cell nuclei. Images are representative of 2 independent experiments. Scale bars 10µm. **(B)** HEK293T or *M. myotis* nasal epithelial cells were infected with WT VSV for 8h at MOI 1. Localization of Dicer (yellow) and dsRNA via J2 antibodies (magenta) were imaged by immunofluorescence and confocal microscopy. DAPI staining (blue) indicates cell nuclei. Images are representative of 2 independent experiments.

